# Divergent evolutionary trajectories shape the postmating transcriptional profiles of conspecific and heterospecifically mated females

**DOI:** 10.1101/2021.11.07.467655

**Authors:** Fernando Diaz, Allan W. Carson, Xingsen Chen, Joshua M. Coleman, Jeremy M. Bono, Luciano M. Matzkin

## Abstract

Postmating-prezygotic (PMPZ) reproductive isolation is hypothesized to result from divergent coevolutionary trajectories of sexual selection and/or sexual conflict in isolated populations (coevolutionary divergence model). However, the genetic basis of PMPZ incompatibilities between species is poorly understood. Here, we use a comparative framework to test predictions of the coevolutionary divergence model using a large transcriptomic dataset generated from con- and heterospecifically mated *Drosophila mojavensis* and *D. arizonae* female reproductive tracts. We found striking divergence between the species in the female postmating transcriptional response to conspecific mating, including differences in differential expression (DE), alternative splicing (AS), and intron retention (IR). As predicted, heterospecific matings produced disrupted transcriptional profiles, but the overall patterns of misregulation were different between the reciprocal crosses. Moreover, we found a positive correlation between postmating transcriptional divergence between species and levels of transcriptional disruption in heterospecific crosses, indicating that mating-responsive genes that have diverged more in expression also have more disrupted transcriptional profiles in heterospecifically mated females. Overall, our results are consistent with predictions of the coevolutionary divergence model and lay the foundation for future studies aimed at identifying specific genes involved in PMPZ incompatibilities and the evolutionary forces that have contributed to their divergence in closely related species.

## Introduction

Speciation results from the accumulation of reproductive isolating barriers that evolve as a consequence of genetic divergence between isolated populations^1^. While historically most speciation research has focused on barriers that arise before mating (premating) or after fertilization (postzygotic), there has been increasing recognition that postmating-prezygotic (PMPZ) isolation is a potent and rapidly evolving barrier to gene flow in many taxa^2,3^. PMPZ isolation arises as a result of incompatible interactions between male and female gametes or among other reproductive molecules involved in postcopulatory interactions. The rapid evolution of PMPZ isolation is consistent with the well-established pattern that genes involved in postcopulatory interactions are among the most rapidly evolving in the genome of many internally fertilizing organisms^4^.

Models of the evolution of PMPZ isolation in animals typically assume that rapid divergence of genes involved in postcopulatory interactions arises as a consequence of divergent trajectories of sexual selection and sexual conflict in isolated populations^2,3^. This is particularly likely in species where females remate frequently because selection on traits mediating male fertilization success and female choice is expected to be intense^5^. Independent coevolutionary dynamics between males and females may follow different paths in diverging populations, leading to alterations in postcopulatory molecular processes that ultimately give rise to PMPZ incompatibilities in crosses between populations^2,3^.

Despite growing evidence highlighting the importance of PMPZ isolation in driving the speciation process, the molecular underpinnings of PMPZ incompatibilities in internally fertilizing animals are not well understood^2^. However, the model based on divergent coevolutionary trajectories of sexual selection and sexual conflict (heretofore the coevolutionary divergence model) makes several predictions regarding the evolution of postmating molecular processes and the genetic basis of PMPZ isolation: (1) Postmating molecular interactions following conspecific mating should diverge between isolated populations as result of different coevolutionary trajectories. (2) Postmating molecular interactions should be disrupted in heterospecific matings. This could include the disruption of postcopulatory processes triggered during the normal response to conspecific mating and/or the induction of processes that are not normally elicited by conspecific mating. (3) Independent coevolutionary trajectories should result in asymmetric patterns of PMPZ isolation in reciprocal crosses, or, in cases where reciprocal crosses both display PMPZ isolation, the molecular basis of incompatibilities is likely to be different. (4) Genes exhibiting more divergence (*e*.*g*. in sequence or transcriptional activity) are expected to be involved in incompatible interactions more than genes exhibiting low levels of divergence.

Although the coevolutionary divergence model predicts rapid divergence in postmating molecular interactions, data comparing postmating responses between closely related species pairs is lacking.However, a recent comparative proteomic analysis of *Drosophila simulans* and *D. mauritiana* revealed extensive species divergence in the proteome of both virgin and mated female reproductive tracts(McCullough *et al*. 2020). Furthermore, only a few studies in plants and animals have explored the molecular basis of PMPZ incompatibilities by comparing postmating transcriptome or proteome responses of con- and heterospecifically-mated female reproductive tissues^6–8^. Heterospecific mating results in disruption of the female postmating transcriptional response in *D. mojavensis and D. novamexicana*, including subsets of genes associated with the normal transcriptional response to conspecific mating, and many additional genes that were not differentially regulated in response to mating in conspecifics^6,7^. In contrast, (McCullough et al. 2020) found few differences in the proteomic response to mating with conspecifics or heterospecifics in *D. simulans* females despite substantial evidence for PMPZ isolation in this cross. Reciprocal crosses were not analyzed in any of these studies, so it is unclear whether the direction of the cross influences the level of disruption. Interestingly, female genes that were transcriptionally misregulated in response to heterospecific mating generally do not evolve more rapidly than other genes in the genome and evolve at lower rates than male seminal fluid protein genes^7,10^. Altogether, results from these studies support some predictions of the coevolutionary divergence model, while others were either not supported or not addressed since experiments included only one direction of heterospecific cross. This highlights the need for additional studies, particularly those where reciprocal crosses can be compared.

In this study, we use the *D. mojavensis/D. arizonae* study system to test predictions of the coevolutionary divergence model in a comparative context. *Drosophila mojavensis* and *D. arizonae* are recently diverged sister species that have long been the focus of speciation research^11^. Previous studies have documented strong PMPZ isolation in crosses involving *D. mojavensis* females, which results in extremely low fertilization success following heterospecific copulation^12^. While the mechanistic basis of these incompatibilities is not fully understood, heterospecifically-mated females exhibit sperm storage defects and fail to efficiently degrade the insemination reaction, which normally forms in the female reproductive tract immediately upon mating^12^. Currently, no data is available to determine the presence or extent of PMPZ isolation in crosses involving *D. arizonae* females. Since evaluation of the coevolutionary divergence model depends on analysis of reciprocal heterospecific crosses, in this study we first demonstrate the presence of strong PMPZ isolation in crosses between *D. arizonae* females and *D. mojavensis* males. We then test predictions of the model using a comparative framework to analyze postmating transcriptomic responses in con- and heterospecifically-mated females for both species. In addition to an analysis of differential expression (DE), we also examine the potential role of alternative splicing (AS) and intron retention (IR) in the postmating transcriptomic response in con- and heterospecifically-mated females. Although comparisons across *Drosophila* show that AS is particularly prevalent in reproductive tissues and contributes significantly to lineage-specific evolution^13^, AS has not been considered in previous studies of the postmating response in female reproductive tissues. We recently demonstrated considerable postmating AS in heads of con- and heterospecifically-mated females^14^, suggesting that AS may play an underappreciated role in the female postmating response. Moreover, disruption of typical patterns of AS may represent an additional mechanism resulting in PMPZ incompatibilities.

## Materials and methods

### Fly stocks

All experiments were carried out using *D. mojavensis* and *D. arizonae* isofemale lines originally collected from Anza Borrego Desert State Park, Borrego Springs, CA (in 2002) and Guaymas, Sonora, Mexico (in 2000), respectively. The genomes of both of these lines have been sequenced and serve as reference sequences for mapping of RNA-seq reads (see below). Flies were held at 25°C, under 12:12 h light:dark cycle and controlled density conditions in 8-dram glass vials with banana-molasses media^15^ for all stocks and experiments.

### PMPZ isolation between D. arizonae females and D. mojavensis males

Previous research has demonstrated strong PMPZ isolation in crosses between *D. mojavensis* females and *D. arizonae* males. To test for PMPZ isolation in the reciprocal cross, we compared fecundity, larval hatching, and fertilization success between con- and heterospecifically mated *D. arizonae* females. Fecundity was analyzed by pairing 8-12 day old virgin *D. arizonae* females with similarly aged either *D. arizonae* or *D. mojavensis* virgin males in an 8-dram glass vial with banana-molasses media. Pairs observed copulating (10-11 pairs per cross) over a 2-hour window were kept and transferred to fresh vials every day for a week. Oviposited eggs were counted, and fecundity was measured as the total amount of oviposited eggs across the seven days. Statistical differences were assessed using a *t*-test. The fecundity vials were used for the determination of hatching rate by incubating them at 25°C for 48 hours and counting total hatched larvae. These data were analyzed with the *afex* package^16^ in R using a generalized linear mixed model (GLMM) with binomial error term and logit link function. Female identity was treated as a random variable to account for the fact that multiple measurements were taken from a single female. Since reduced larval hatching could result from embryo inviability or lack of fertilization, we performed an additional experiment to help differentiate these possibilities. Virgin 8-12 old *D. arizonae* females were paired in vials with either a *D. arizonae* or *D. mojavensis* virgin male (8-12 days old). Copulations were observed (N=60) and mated females were held together overnight in a large population cage to oviposit. In the morning, the plate was removed and stored for 6 hrs to ensure any developing embryos would be six to 22 hrs old. We then stained eggs with 4′,6-diamidino-2-phenylindole (DAPI) to determine whether embryonic development had been initiated. We considered eggs to be unfertilized if there was no indication that development had commenced and there were no more than four DAPI stained nuclei present (representing four products of female meiosis). Fertilization data was analyzed in R using a generalized linear model (GLM) with binomial error term and logit link function. To evaluate sperm storage, we conspecifically and heterospecifically paired up virgin *D. arizonae* and *D. mojavensis* females as described for the fecundity assay. Following successful copulation, males were removed, and females were pooled in vials (2-10 females per vials). Females were either maintained at 25°C for 24 hours or five days before dissection. Female LRTs were dissected in sperm buffer (0.05M Tris, 1.1% NaCl, 0.1% dextrose, 0.01% L-arginine, 0.01% L-lysine), placed on a slide, covered with a glass coverslip and visualized on an Olympus inverted light microscope. Sperm motility within the seminal receptacle was classified into one of four categories: many (wave-like sperm tail activity observed across nearly the entirety of the seminal receptacle); medium (one or two regions of wave-like sperm tail motility); few (tens or fewer visible motile sperm scattered along the seminal receptacle lumen); none (no motile sperm). Motility categories were analyzed as ordinal variables using an ordinal logistic regression in JMP v16.

### Experimental design used for differential gene expression and alternative splicing

Experimental design consisted of conspecific and heterospecific matings between the species. We refer to the reciprocal mating as ♀*Dmoj* for con- and heterospecific matings involving *D. mojavensis* female, and ♀*Dari* for those involving *D. arizonae* females. All mating experiments were performed using virgin flies (8-12 days old) by isolating pairs in vials and observing copulation events during a 2-hour window in the morning. Males were removed from vials after copulation and females were kept in the vials until the specified postmating period was reached (45 min or 6 hrs), when lower reproductive tracts (LRTs) were collected from both mated and virgin females. Groups of 20 specimens were pooled for each sample and three replicates were collected per experimental cross, which generated a total of 30 samples. All tissues were placed immediately in TRIzol and kept at -80 °C until total RNA extractions.

### RNA extraction, cDNA library construction, and sequencing

Total RNA was extracted using Direct-zol RNA kit (Zymo Research). Both RNA quality and quantity were inspected on a Bioanalyzer (Applied Biosystems/Ambion). cDNA libraries were generated using KAPA Stranded mRNA-Seq Kit following manufacturer’s instructions. Libraries were sequenced at Novogene Inc. using the HiSeq SBS v4 High Output Kit on Illumina platform flow cells with runs of 2 × 150 bp paired-end reads. Illumina’s HiSeq Control Software and CASAVA software (Illumina, Inc.) were used for base calling and sample demultiplexing.

### Sequence trimming and mapping

Nearly 700 million total paired-end read sequences were obtained from the Illumina runs, ranging from 11 to 34 million raw paired-end reads for each sample. Reads were trimmed for quality and adapter sequences were removed using a minimum quality base of Q = 20 and minimum read length of 50 bp using the software Trimmomatic^17^. Trimmed reads were then mapped to corresponding reference genomes using splice-aware mapper *GSNAP*^18^ with the option of new splice events detection. Template based genomes from the same *D. mojavensis* and *D. arizonae* lines utilized in this study were used for mapping RNAseq reads. For *D. mojavensis*, the assembly from ^19^ (Accession number SRP190536) was used with updated annotations retrieved from FlyBase version FB2016_05^20^. A template genome version of *D. arizonae* (osf.io/ukexv) was assembled using the same method as *D. mojavensis* in ^19^ with paired-end and mate pair Illumina reads from^21^ (Accession number SRP278895). Generated *sam* files were converted to *bam* format after indexing and filtering for a minimum mapping quality of MQ = 20 using SAMtools ^22^. These mapping results were then used for all differential expression and alternative splicing downstream pipelines. There is previous evidence suggesting the transfer of male accessory gland-derived transcripts to the females LRT during copulation when crossing *D. arizonae* males to *D mojavensis* females^6^. Therefore, to ensure that our transcriptional analysis corresponded to the female transcriptional response, we removed all genes whose transcripts were completely male-derived from subsequent analyses. Any transcripts with sequencing reads from both males and females were retained in the dataset.

### Differential expression (DE)

We created a gene level read count matrix for all samples using *featureCounts*^23^. The read count matrix was filtered for a minimum count cutoff of 3 cpm in at least two out of three replicates per group. All DE analyses were performed using the R package *edgeR*^24^ after *TMM* library normalization. Normalized counts were analyzed by a GLM accounting for negative binomial variable of read counts, followed by DE analyses. All comparisons were performed between mated females (con- and heterospecific mating) and virgin females (♀*Dmoj* and ♀*Dari*) at each postmating period (45 min and 6 hrs). An *FDR* correction following a global α of 0.05 was used^25^ for multiple comparisons, as well as a log_2_-fold-change threshold of 1.0.

### Alternative splicing (AS) and intron retention (IR)

We used *JunctionSeq*^26^ to detect genome-wide patterns of alternatively spliced genes. The pipeline is based on differential usage calculated from both exon and junction feature coverages. A new flattened *GTF* annotation file that excluded overlapping features was first generated using *QoRTs*^27^. All overlapping genes were merged as composed by a flat set of non-overlapping exons and splice junctions with unique identifiers. *QoRTs* was also used to generate a read count matrix for AS analysis, including three types of read counts per gene as estimated by exons, junction and gene level counts. No read was counted more than once in the model since exon and junction dispersions are fitted independently. As for DE analysis, alternatively spliced genes were detected if at least one exon or splice junction was differentially used (relative to overall expression of the gene) between mated females (con- and heterospecific mating) and virgin females (♀*Dmoj* and ♀*Dari*) at each postmating period (45 min and 6h). An *FDR* correction following a global α of 0.01 was used^25^ for multiple comparisons, as well as a log_2_-fold-change threshold of 1.0.

Intron retention is a specific type of AS that is not necessarily captured by *JunctionSeq* and is generally considered as a mechanism of expression downregulation, as transcripts with retained introns are degraded by the nonsense-mediated decay pathway^28^. However, in some cases transcripts with retained introns have been shown to perform novel functions^29–31^. We investigated whether postmating AS events also involve mechanisms of intron retention using the *IRFinder* pipeline^32^. For each genome, a new reference annotation was built by removing all overlapping features present of individual introns and then unique identifiers were assigned to each flattened exon. Only regions with high mapping scores as estimated through simulated reads across the genome were included in the flattened annotation file. A read count matrix with all reads overlapping splice junctions was generated and IR rates were estimated as junction reads / (junction reads + intronic reads) for each sample. The count matrix was then used by *IRFinder* R package^32^ in order to estimate the GLM. This method was used to test the fold change of IR between biological conditions using the *DESeq2* R package framework^33^. As for differential expression analysis, alternatively spliced genes were then detected if least one exon or splice junction was differentially used between mated females (con- and heterospecific mating) and virgin females (♀*Dmoj* and ♀*Dari*) at each postmating period (45 min and 6h). An FDR correction following a global α of 0.05 was used^25^ for multiple comparisons.

Because IR changes could serve as a mechanism of downregulation by transcript degradation, we tested this hypothesis by estimating IR changes between mated vs virgin samples (IR change), while comparing up vs down-regulated genes. A GLM analysis was performed using categories of up and down regulation as independent variables and the level of IR change as the dependent variable for each mating experiment. GLM analysis was performed using a gaussian family distribution with identity link function after square root transformation of data to ensure assumptions of the GLM were met.

### Postmating transcriptional divergence and disruption of gene expression in heterospecific matings

To investigate whether genes that have diverged in expression between the species are more likely to be disrupted by heterospecific matings, we compared the proportion of misregulated genes in three categories: (1) genes DE in conspecific *D. mojavensis* matings but not conspecific *D. arizonae* matings, (2) genes DE in conspecific *D. arizonae* matings but not conspecific *D. mojavensis* matings, and (3) genes DE in both conspecific crosses. Differential expression data was analyzed using a GLM in R with a binomial response variable and logit link function. The full factorial model included the factors gene category, postmating timepoint, and the interaction. For the analysis of AS data, the model did not include time due to the small number of genes in some categories at the 45 min timepoint. *Post-hoc* testing using Tukey’s adjustment was conducted with the R package *emmeans* (Lenth).

We then investigated whether the level of heterospecific expression disruption can be predicted by transcriptional divergence between the species. For this, we estimated the level of heterospecific disruption as the Euclidian distance of relative expression values (fold change relative to virgin) between hetero-vs conspecific transcriptional responses (heterospecific disruption = heterospecific response - conspecific response). Postmating transcriptional divergence between the species was estimated as the Euclidean distance of conspecific relative expression values (*e*.*g*., *D. mojavensis* conspecific expression divergence = conspecific *D. arizonae* - conspecific *D. mojavensis*). Transcriptional correlations were tested based on Pearson’s correlation coefficients for all genes differentially regulated in con- or heterospecific crosses, and separately for all conspecific mating responsive genes (including those also differentially regulated in the heterospecific cross), and genes only differentially expressed in response to a heterospecific mating.

### Functional and molecular evolutionary analyses

Overrepresentation of specific categories of biological process were then investigated for DE and AS genes using *Panther*^35^. Additionally, we investigated signatures of positive selection on genes responding to con- vs heterospecific matings. For this, we estimated evolutionary rates (ω = *d*_*n*_*/d*_*s*_) using *codeml*, part of *PAML* 4.9^36^. *CDS* alignments between *D. mojavensis* and *D. arizonae* were produced with *MUSCLE* 3.8.31^37^. Any alignments with internal stop codons or frameshifts were removed before analysis. *Codeml* was run using model *0* with default values. Raw synonymous and nonsynonymous polymorphism counts were generated with *KaKs* Calculator 1.2^38^ and loci lacking synonymous substitutions were omitted from the analysis. Pooled gene lists from both time points were used for all statistical analyses. For each species and cross type, significant deviation from the genome-wide ω was determined using the Dunn method for joint ranking.

## Results

### Postmating-prezygotic isolation is strong in crosses between D. arizonae females and D. mojavensis males

Heterospecifically-mated *D. arizonae* females laid 23% fewer eggs over the course of seven days postmating compared to conspecifically mated females (t-test, t=2.15, P = 0.04; Fig 1a) and a much smaller proportion of eggs laid by heterospecifically mated females hatched (GLMM, X^2^=26.9, P < 0.001; Fig 1b). Analysis of DAPI stained eggs indicated that most unhatched eggs laid by heterospecifically mated females were unfertilized (GLM, X^2^=242.9, P < 0.001; Fig 1c). The effect of heterospecific mating on sperm storage was dramatic in *D. arizonae* females where the many and medium motility categories fell from ∼85% in conspecifically mated females to ∼13% in heterospecifically mated females at both 1 and 5 days post-mating (X^2^=17.8, P < 0.001 and X^2^=20.1, P < 0.001, respectively; Fig 1d). A similar decline in sperm motility was observed in *D. mojavensis* at 5 days post-mating (X^2^=16.6, P < 0.001), but no significance was observed at one day (Fig 1d).

**Fig. 1.**
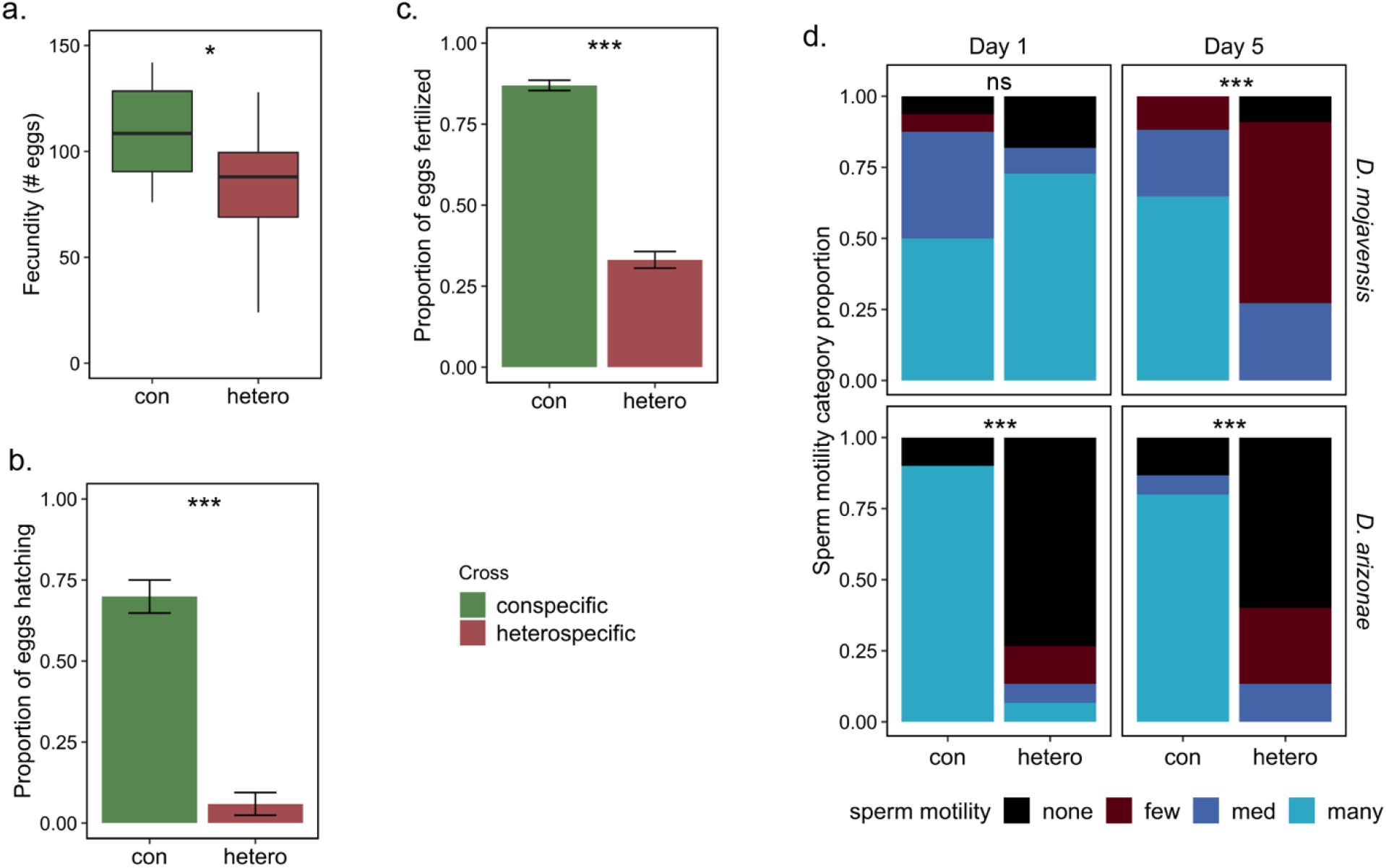
Evidence of postmating-prezygotic isolation between *D. mojavensis* and *D. arizonae*. **a**. Egg production over seven days of *D. arizonae* females following a con- or heterospecific mating. **b**. Proportion of eggs oviposited by *D. arizonae* females in **a**., that hatch into first instar larva. **c**. Proportion of eggs oviposited by *D. arizonae* females following a con- or heterospecific mating that were determined to be fertilized using DAPI staining. **d**. Levels of sperm motility within seminal receptacles in both *D. mojavensis* and *D. arizonae* females mated con- and heterospecifically at 1 and 5 days postmating. P-values of correlations are noted: * P < 0.05, ** P < 0.01, *** P < 0.001, ns not significant.

### Transcriptional regulation in response to mating includes multiple mechanisms

We obtained an average of 20*10^6^ mapped reads for each library following trimming and filtering of sequence reads. Minimum count filtering was applied independently to all gene subfeatures at the beginning of each analysis (*e*.*g*., exon, junction, intron). We found evidence for significant transcriptional changes when comparing mated vs virgin samples (Fig 2a). These changes were detected at all types of gene regulation analyzed including DE, AS, and IR (Fig 2a). Few genes were found to be differentially regulated by multiple mechanisms, indicating that DE, AS, and IR represent distinct postmating responses that target different genes (Fig 2a). In accordance with predictions based on the known role of IR as a mechanism of expression downregulation, we found that genes with higher intron retention were significantly downregulated compared to genes with lower rates of intron retention (Fig S1).

**Fig. 2.**
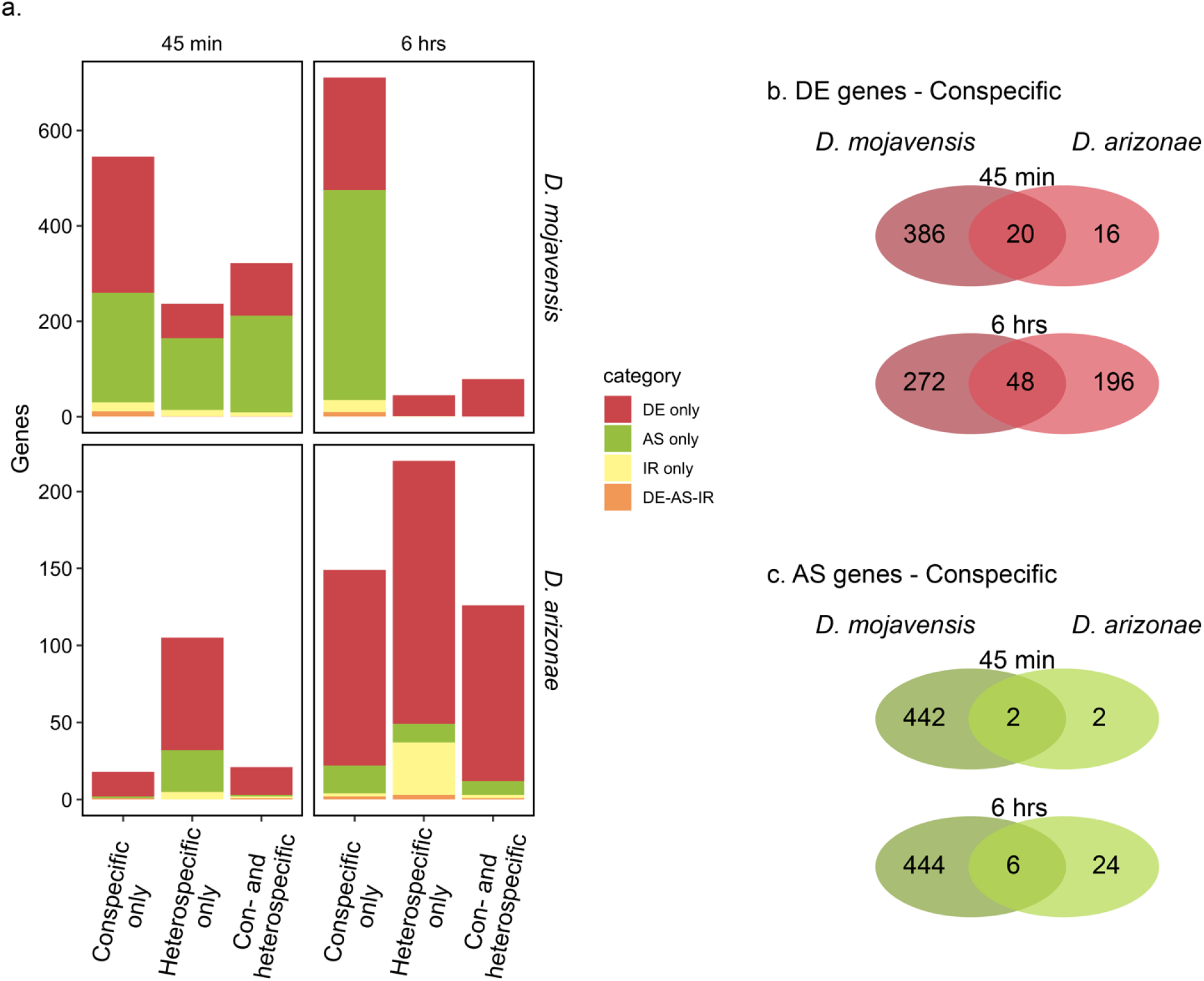
Patterns of transcriptional and splicing variation. **a**. Number of genes with significant patterns of DE, AS and IR following con- and heterospecific matings between *D. mojavensis* and *D. arizonae*. All comparisons were performed against virgin females at 45 min and 6 hrs postmating (*FDR*_*α*_ = 0.05). Genes in the DE-AS-IR category, show significance in two or more of the individual categories (DE, AS and/or IR). The bars represent the number of genes exclusive to con- or heterospecific matings (Conspecific or Heterospecific only), as well as the number of overlapping genes (Common to Con- and Heterospecifics). **b**. Comparison of the number of genes differentially expressed in response to a conspecific mating within and between each species. **c**. Comparison of the number of genes alternatively spliced in response to a conspecific mating within and between each species.

### Transcriptional response to mating has diverged between D. mojavensis and D. arizonae

Quantitative transcriptional changes for genes that responded to mating in conspecific crosses was significantly positively correlated between the species (*R*^*2*^ = 0.23, P < 0.001 and *R*^*2*^ = 0.12, P < 0.001, at 45 min and 6 hrs respectively; Fig 3). Moreover, the correlation for mating responsive genes was much higher than that for non-mating responsive genes (Fig S2). While together these results indicate some conservation of the postmating transcriptional response between *D. mojavensis* and *D. arizonae*, most of the variation in the postmating response (77% and 88% at 45 min and 6 hrs respectively) was not explained by the regression model and many genes displayed highly discordant patterns of expression between the two species (e.g. those in quadrants II and IV of Fig 3). Consistent with this, comparison of DE and AS between the species revealed little overlap in the genes that were differentially regulated in response to mating (Fig 2b). Moreover, the overall strength and temporal pattern of transcriptional response varied between the species. *Drosophila mojavensis* displayed a more rapid and stronger transcriptional response to mating that included substantial DE and AS. In contrast, in *D. arizonae* there was a much weaker initial transcriptional response to mating that increased over time. Moreover, fewer genes were differentially expressed, and the amount of alternative splicing was significantly reduced compared to *D. mojavensis*.

**Fig. 3.**
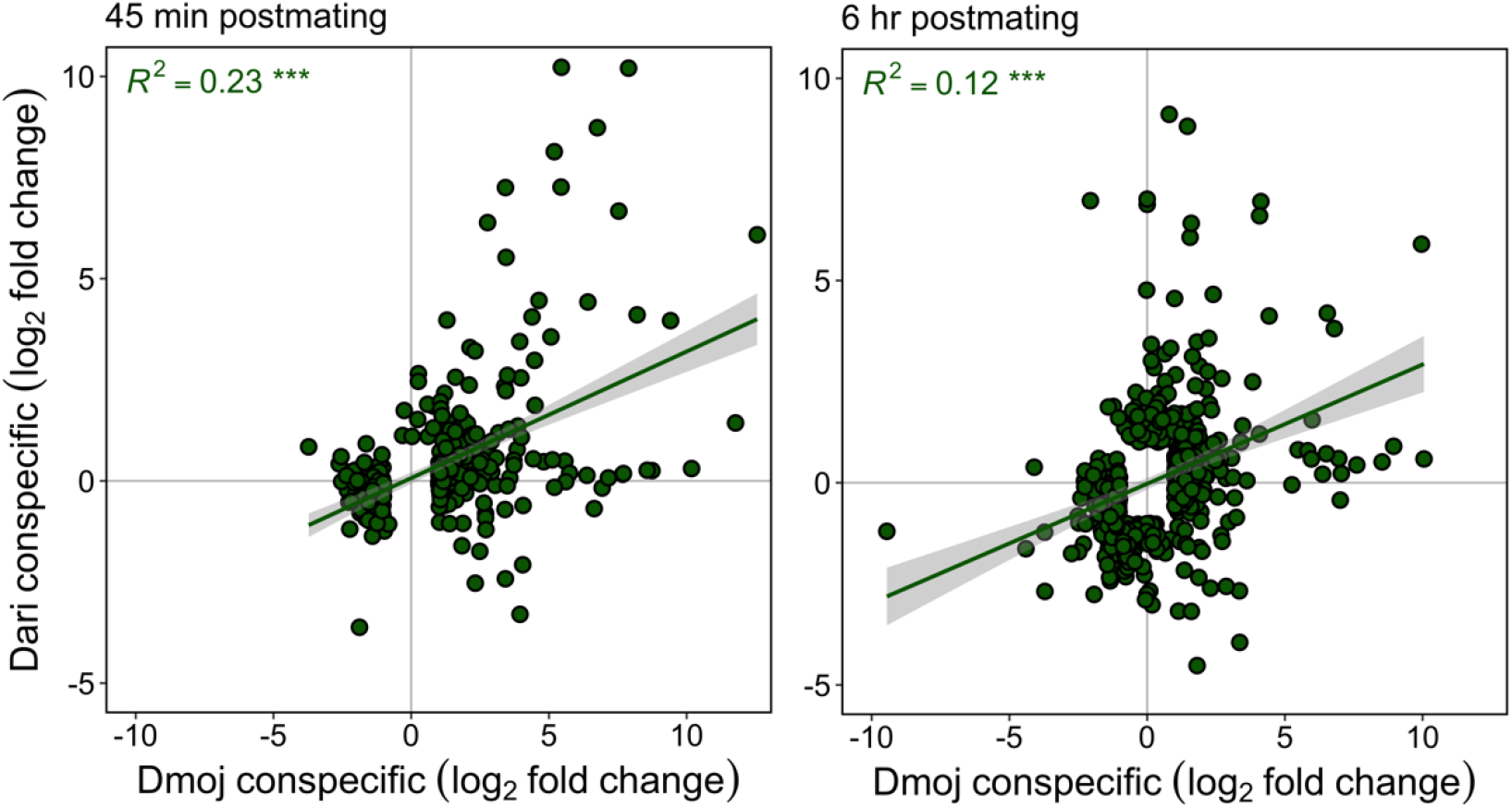
Transcriptional correlations between conspecific-responsive DE genes in *D. mojavensis* vs *D. arizonae*. All comparisons are performed against virgin females at 45 min and 6 hrs postmating (*FDR*_*α*_ = 0.05). Scatterplots represent the relationship of relative fold changes (log_2_) for conspecific matings in *D. mojavensis* vs *D. arizonae*. Pearson’s *R*^*2*^ correlation coefficients and LM trend-lines (with 95% confidence intervals) between the species are indicated for conspecific-responsive genes expressed in *Dmoj, Dari* and overlapping genes responding in both species. P-values of correlations are noted: *** P < 0.001.

GO-term enrichment analysis of DE genes revealed several overrepresented terms in conspecific crosses (Fig 4). There were fewer enriched terms in *D. arizonae* than *D. mojavensis*, which likely reflects the overall smaller number of DE genes in this species. None of the enriched terms overlapped between the species, potentially indicating functional divergence in the postmating transcriptional response. GO-term enrichment analysis of AS genes did not detect any overrepresented categories in either conspecific cross.

**Fig. 4.**
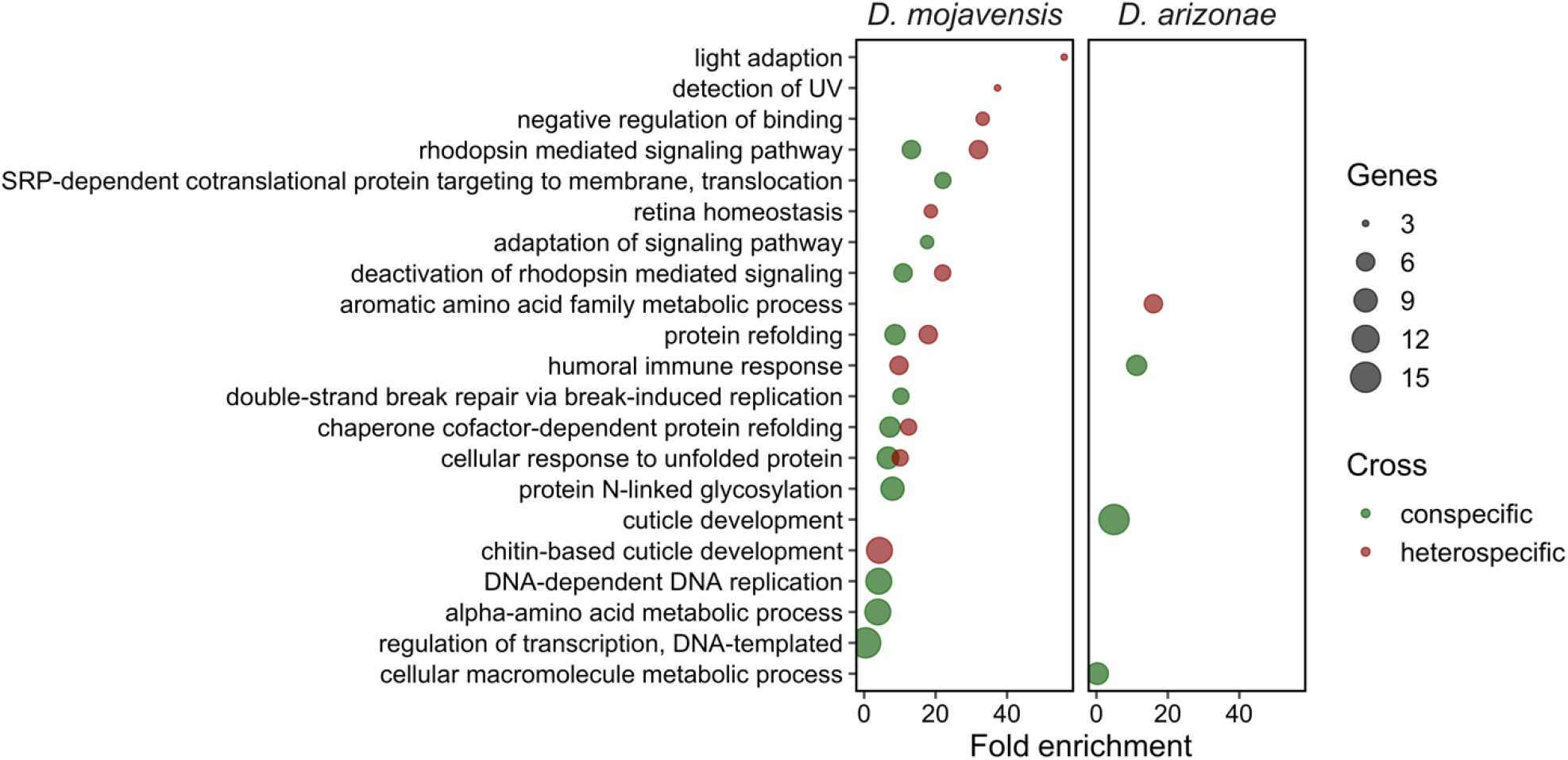
Gene ontology analysis of DE genes. Functional analysis for DE genes, indicating biological process gene ontology enrichment categories for con- vs heterospecific matings. The fold enrichment of detected genes within each enriched category is indicated.

### Transcriptional response to mating is disrupted in heterospecific crosses

The number of genes that were differentially regulated (DE and AS) in both con-and heterospecific crosses was relatively low (Fig 2), indicating the overall transcriptional response to mating in heterospecifically mated females was distinct form the conspecific response. In addition, there were a relatively large number of genes, particularly in *D. arizonae*, that were differentially regulated only in the heterospecific cross (Fig 2). GO-term enrichment analysis of DE genes in heterospecific crosses overlapped to some degree with conspecific crosses, although a number of terms were enriched in one cross but not the other, indicating that transcriptional disruptions in heterospecific crosses could have functional consequences (Fig 4).

In *D. mojavensis*, although ∼75% of conspecific mating responsive genes exhibited disrupted expression in heterospecifically mated females at 45 min postmating (Fig 5), the overall pattern of response between the crosses was highly correlated (*R*^*2*^ = 0.83, P < 0.001; Fig 6a). This indicates that most transcriptional disruptions were of small magnitude early on after mating. At 6 hrs the proportion of genes with disrupted expression profiles remained high, and the magnitude of the differences also increased (*R*^*2*^ = 0.41, P < 0.001; Fig 6a). In *D. arizonae*, most conspecific mating responsive genes also exhibited disrupted expression profiles (Fig 5). However, in this case the magnitude of expression differences was greater at 45 min than 6 hrs (*R*^*2*^ = 0.38, P < 0.001 and *R*^*2*^ = 0.58, P < 0.001, respectively; Fig 6b). Across each timepoint, genes that were differentially regulated in both conspecific crosses were less likely to show disrupted expression profiles in heterospecific matings than genes that were only differentially regulated in one species (GLM, *X*^*2*^=4.1, P= 0.100 for category × time interaction; *X*^*2*^=58.7, P < 0.001 for category; *post-hoc* tests for gene category: both – moj, z-ratio= -5.9, P < 0.001; ari – both, z-ratio= 2.6, P= 0.026; Fig 5). This serves to underscore the fact that while the transcriptional response to mating in both species is disrupted, the pattern of disruption is largely unique to each species. No such relationship was found for alternative splicing as there were no differences between the gene categories (*GLM, X*^*2*^=1.4, P= 0.496).

**Fig. 5.**
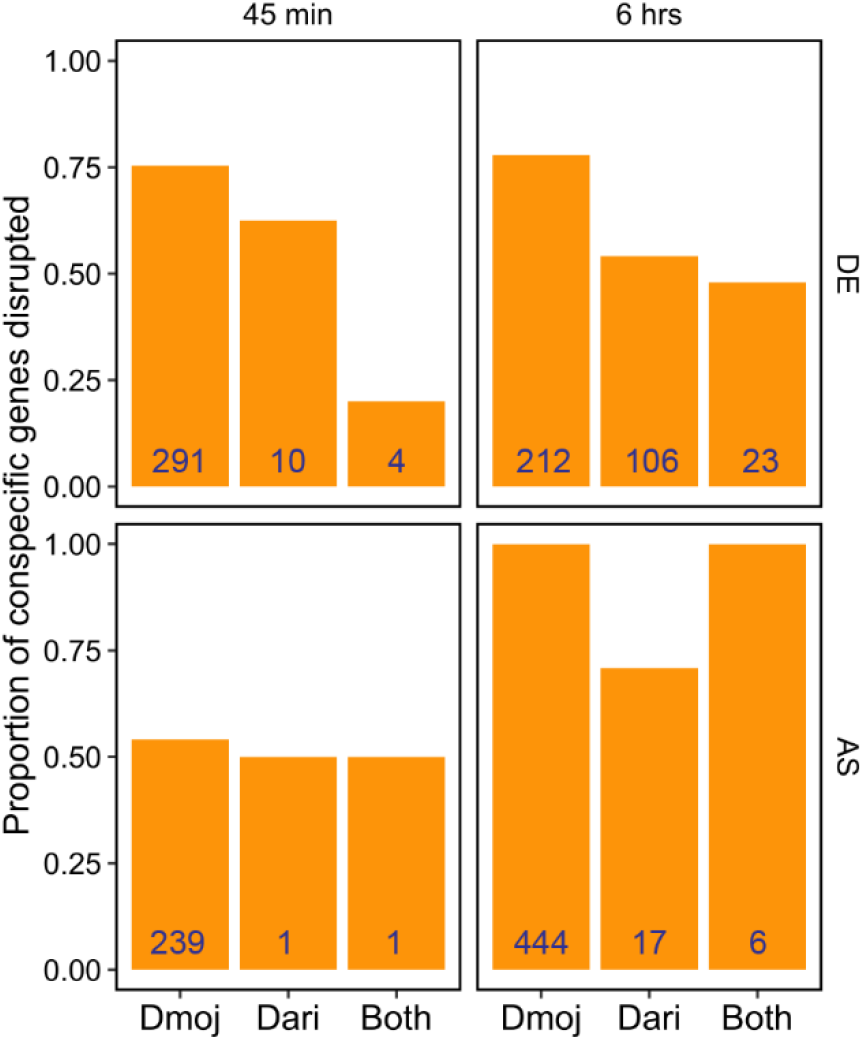
Heterospecific disruption rates for conspecific-mating responsive genes. The proportion of conspecific-mating responsive genes experiencing heterospecific disruption at 45 min and 6 hrs postmating (*FDR*_*α*_ = 0.05). The bars represent the number of differentially regulated genes exclusive to *D. mojavensis* (Dmoj) or *D. arizonae* (Dari), and genes that were differentially regulated in both species (Both). See text for statistical analysis. Numbers in blue within bars indicate the number of genes with disrupted expression profiles.

**Fig. 6.**
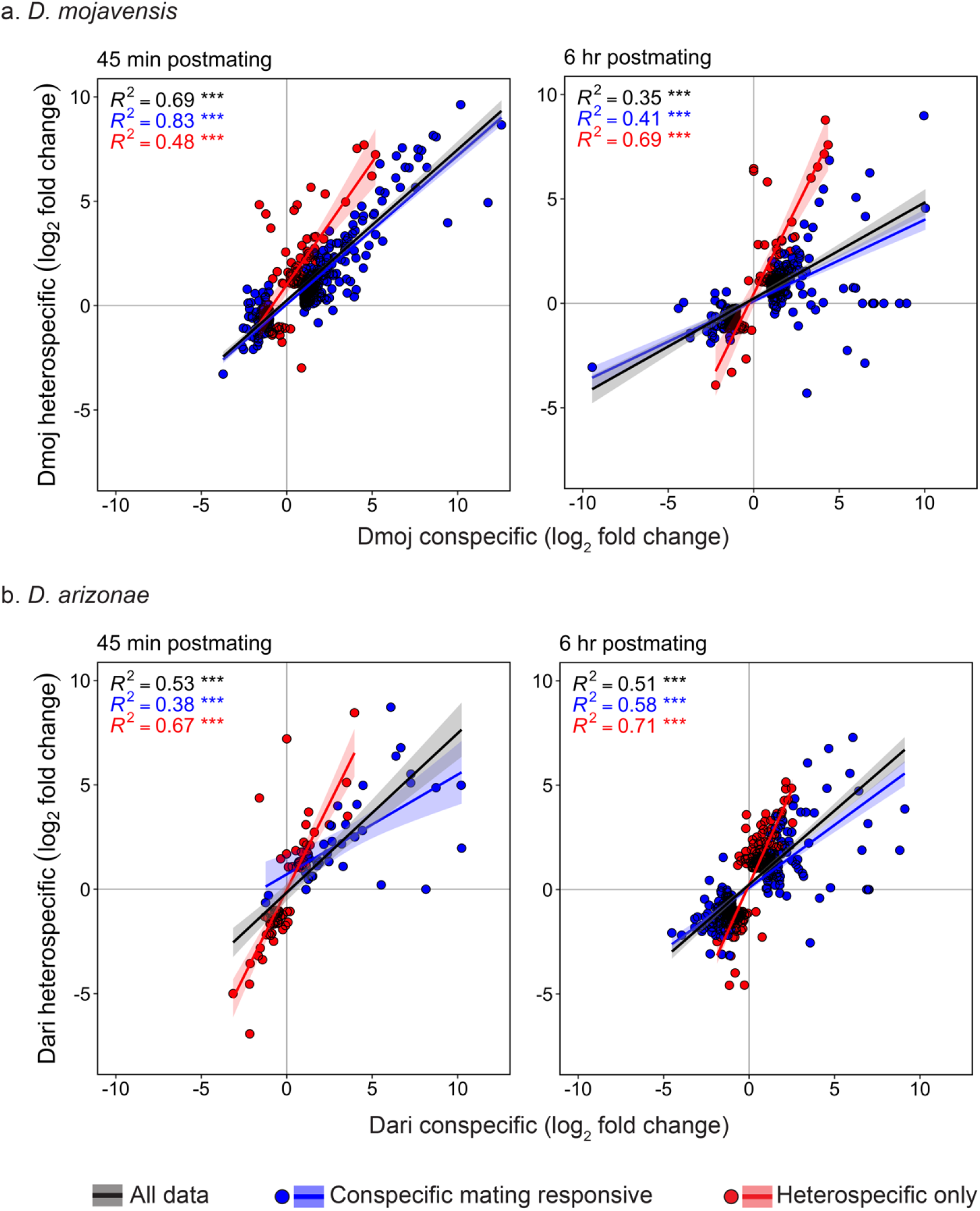
Transcriptional correlations for DE genes between con- vs heterospecific matings in *D. mojavensis* and *D. arizonae*. All comparisons were performed against virgin females (*FDR*_*α*_ = 0.05) for **a**. *D. mojavensis* and **b**. *D. arizonae* matings. Scatterplots represent the relationship of relative fold changes (log_2_) between con- vs heterospecific matings at 45 min and 6 hrs postmating. Colors indicate the expression change of genes showing a conspecific mating response (conspecific only + conspecific/heterospecific overlap) or heterospecific only response (blue and red points respectively), as well as all the genes (black points). Pairwise Pearson’s *R*^*2*^ correlation coefficients and LM trend-lines (with 95% confidence intervals) between con- vs heterospecific crosses are indicated for con-, heterospecific and overlapping genes responding in both crosses. P-values of correlations are noted: *** P < 0.001.

### Transcriptional divergence between species predicts transcriptional disruption in heterospecific crosses

In *D. mojavensis*, the magnitude of expression disruption in heterospecific crosses was significantly positively correlated with expression divergence between the species (Fig 7a). As the magnitude of expression disruption increased over time (Fig 6a) for conspecific mating responsive genes, the correlation between disruption and divergence strengthened as well (*R*^*2*^ = 0.42, P < 0.001 and *R*^*2*^ = 0.70, P < 0.001, for 45 min and 6hrs respectively; Fig 7a). Likewise, there was a positive correlation between disruption and divergence in *D. arizonae*, though the strength of the correlation was similar at both time points (Fig 7b). Notably, in both species these positive correlations were driven by the pattern in conspecific mating-responsive genes. There was either no correlation or a slight negative correlation between disruption and divergence in genes that were differentially regulated only in heterospecific crosses.

**Fig. 7.**
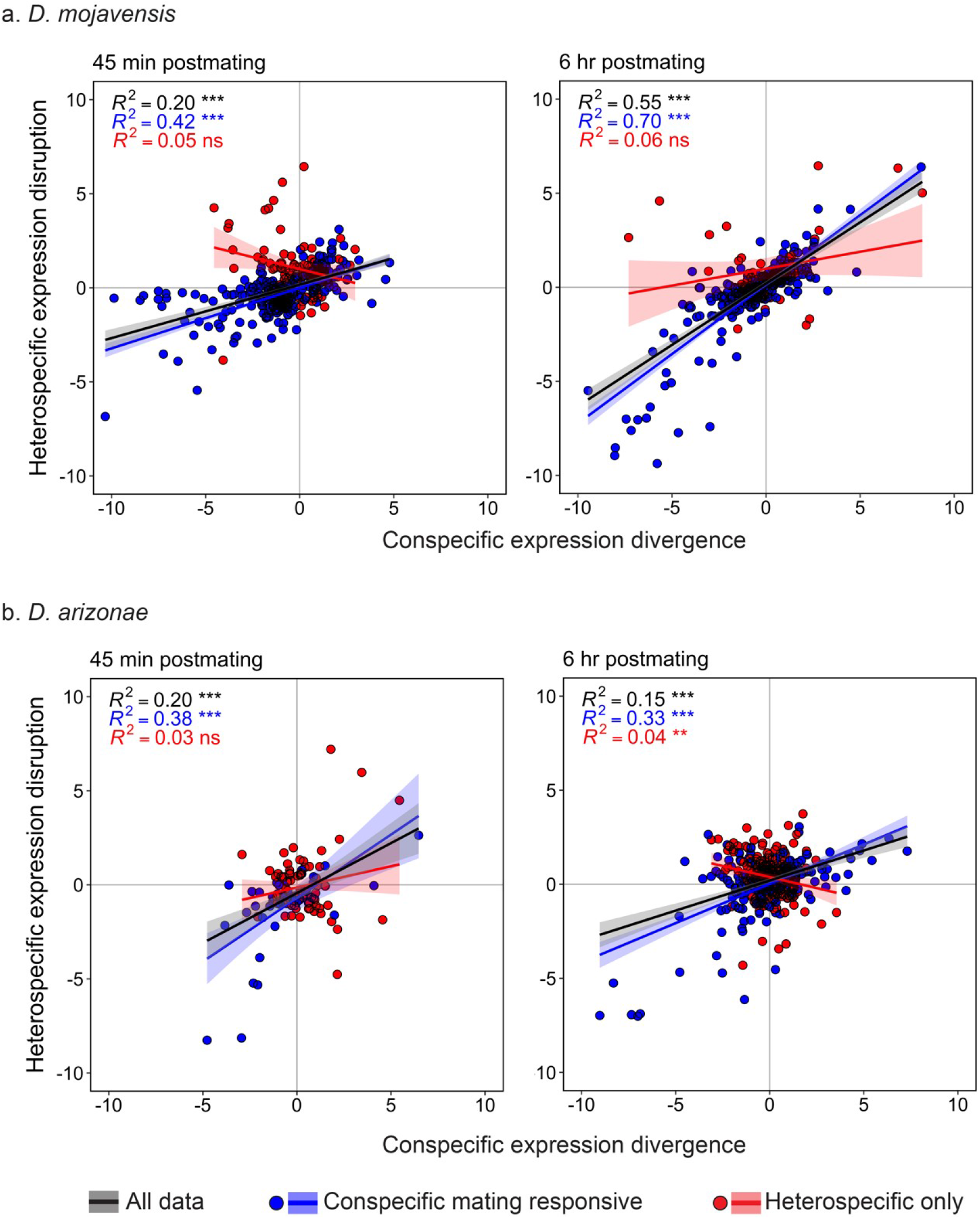
Relationship between postmating conspecific expression divergence and the level of heterospecific disruption for conspecific-responsive DE genes. Scatterplots show the relationship between postmating conspecific divergence (*e*.*g*., *D. mojavensis* conspecific expression divergence = conspecific *D. arizonae* - conspecific *D. mojavensis*) and the level of heterospecific disruption (*e*.*g*., heterospecific disruption in *D. mojavensis* = heterospecific *D. mojavensis* - conspecific *D. mojavensis*). Only DE genes detected in con- or heterospecific at 45 min and 6 hrs postmating are considered (*FDR*_*α*_ = 0.05). Colors indicate the expression change of genes showing a conspecific mating response (conspecific only + conspecific/heterospecific overlap) or heterospecific only response (blue and red points respectively), as well as all the genes (black points). Pairwise Pearson’s *R*^*2*^ correlation coefficients and LM trend-lines (with 95% confidence intervals) between con-vs heterospecific crosses are indicated for con-, heterospecific and overlapping genes responding in both crosses. P-values of correlations are noted: ** P < 0.01, *** P < 0.001, ns not significant.

In *D. mojavensis*, genes with disrupted expression profiles (DE or AS), including conspecific mating responsive genes and those that were differentially regulated only in heterospecific crosses, exhibited lower ω values compared to the genome background in three out of four comparisons (z = -3.86, P < 0.001 for conspecific DE; z = -8.74, P < 0.001 and z = -4.11, P < 0.001 for con- and heterospecific AS, respectively; Fig 8). In contrast, there were no significant differences between genes with disrupted expression profiles and the genome background in *D. arizonae* (Fig. 8).

**Fig 8.**
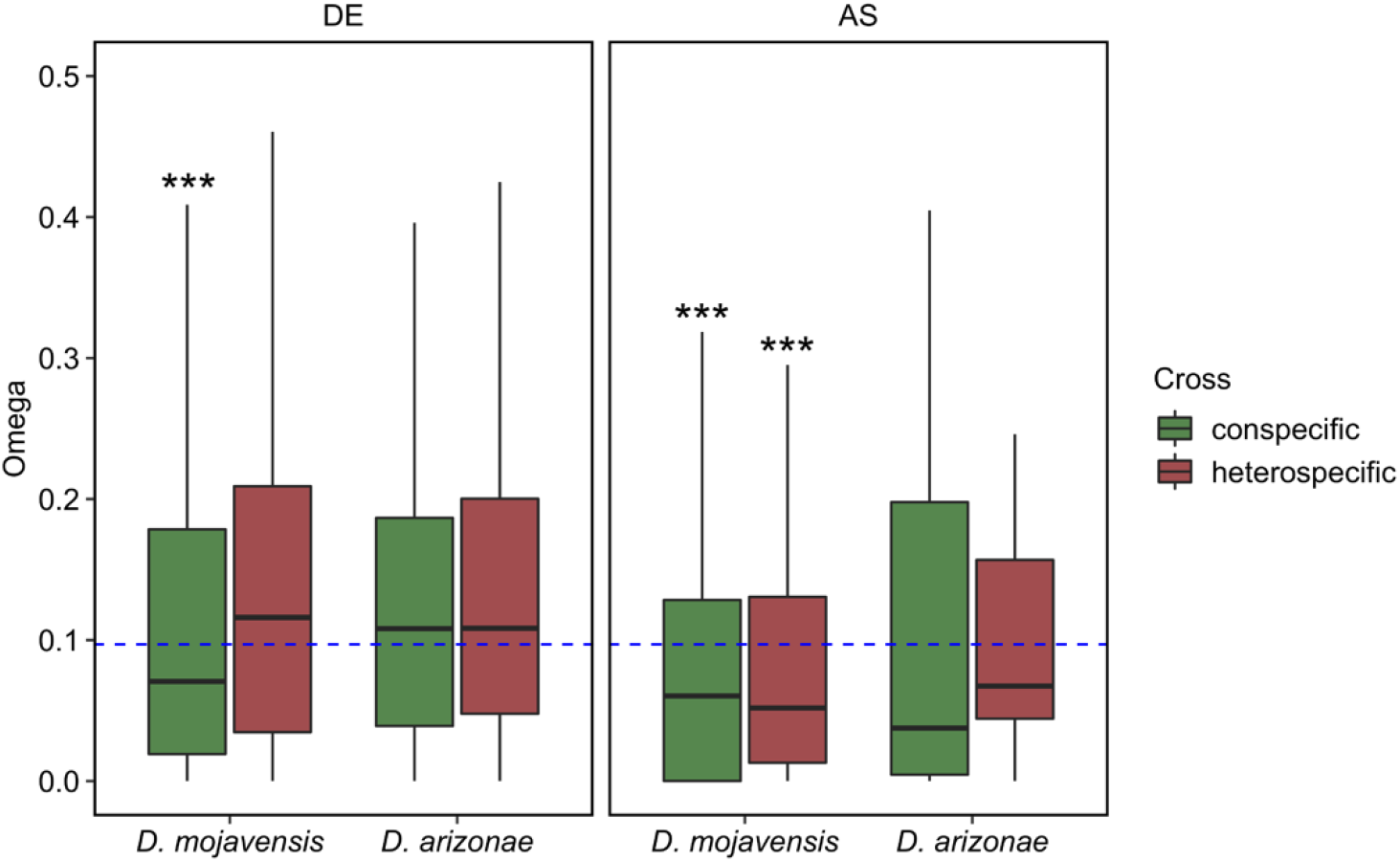
Average pairwise dN/dS (ω) for DE and AS genes induced by con- and heterospecific matings. The median ω for the genome background (blue dash line) is indicated. Significant comparisons against the genome background using the Dunn method for joint ranking are indicated. P-values are noted: *** P < 0.001.

## Discussion

Previous studies have shown that PMPZ barriers in crosses between all four cactus host populations^39^ of *D. mojavensis* females and *D. arizonae* males are strong^12^. Here, we demonstrate PMPZ isolation in crosses between *D. arizonae* females and Mojave *D. mojavensis* males, which further underscores the importance of reproductive tract incompatibilities in this system and provides a comparative framework for examining the molecular basis of PMPZ incompatibilities. The reduction in fecundity we observed for *D. arizonae* females mated to *D. mojavensis* males is largely consistent with a previous study showing similar reductions oviposition by heterospecifically mated *D. mojavensis* females from three of the four cactus host populations^12^. Notably, the one exception to this pattern was found in the reciprocal cross using *D. mojavensis* from the Mojave Desert population (as in this study), which laid more eggs when mated to heterospecifics than conspecifics. The difference between the reciprocal crosses suggests divergence in the mechanisms driving PMPZ isolation. While both heterospecific crosses exhibit severe reductions in fertilization success, the underlying incompatibilities that result in fertilization failure are not well understood. A previous study demonstrated that reduced sperm viability within the female reproductive tract likely contributes to a reduction in fertilization success in crosses between *D. mojavensis* females and *D. arizonae* males^12^. Although we observed similar results in *D. mojavensis*, we also found that crosses between *D. arizonae* females and *D. mojavensis* males were even more severely affected. Specifically, at one day postmating the majority of heterospecific sperm in the seminal receptacle of *D. mojavensis* females was motile, while the presence of motile sperm was sharply reduced at five days postmating. In contrast, in *D. arizonae* we detected few motile heterospecific sperm even at one day postmating. Although these data suggest that sperm inviability in the female reproductive tract likely contributes to PMPZ isolation in both crosses, the underlying incompatibilities that affect sperm viability may not be the same given differences in the severity and time course of the observed phenotypes. It is likely that other factors also contribute to the reduction in fertilization success, particularly in *D. mojavensis* where females produce few fertilized eggs even during the time period where heterospecific sperm are still motile. Overall, these findings further establish the utility and power of the *D. mojavensis/D. arizonae* study system for understanding the evolution and mechanistic basis of PMPZ isolation, including testing predictions generated from the coevolutionary divergence model.

The coevolutionary divergence model predicts that species pairs, such as *D. mojavensis* and *D. arizonae*, should exhibit divergent transcriptional responses to conspecific mating as a result of independent coevolutionary trajectories between males and females of each species. As expected for recently diverged species, there was a positive correlation between the transcriptomic response to conspecific mating between the two species, which declined from 45 min to 6 hrs postmating. Nevertheless, these correlations were relatively weak, indicating substantial divergence between the species in the overall transcriptional response to mating. This conclusion is further supported by fundamental differences in transcriptional patterns of DE, AS, and functional categories associated with mating responsive genes in each species.

In general, *D. mojavensis* females displayed a more comprehensive transcriptomic response to mating than *D. arizonae* females, especially at the 45 min postmating timepoint where nearly 11-fold more genes were DE in *D. mojavensis* females. This difference persisted at 6 hrs postmating though the magnitude of the difference was smaller. While *D. mojavensis* showed a stronger early transcriptional response that declined over time, *D. arizonae* displayed the opposite pattern, with more DE genes detected at 6 hrs than at 45 min. Interestingly, only 5% of all mating responsive DE genes were differentially regulated in both species, demonstrating striking divergence in the overall pattern of differential expression. Moreover, GO-term enrichment analysis of DE genes suggests potential functional divergence in the postmating response as we found nearly no overlapping enriched categories between the species.

Alternative splicing is known to be an important form of gene regulation in *Drosophila* female reproductive tissues, but, to our knowledge, has not been examined in the context of the postmating transcriptional response to mating. We found that AS is a key component of the female postmating response and that alternatively spliced genes are largely distinct from genes that are differentially expressed following mating (less than 5% of genes were found in both categories). Together with our recent report of significant alternative splicing regulation in the head following mating in these species^14^, our results suggest AS is an important but previously overlooked mechanism of the female transcriptomic response to mating. While most of the AS detected likely affects regulation of different protein isoforms, we also found differences in intron retention for many genes. Intron retention is a known mechanism of downregulation of gene expression as transcripts with retained introns trigger the nonsense mediated decay (NMD) pathway^28^. Consistent with this, we found intron retention was associated with transcriptional downregulation of genes following mating, which also concurs with our previous findings of postmating transcriptional responses in the female head^14^. Alternative splicing analysis also revealed substantial differences between species in the overall role of AS in the postmating response. Like the DE analysis, we found that AS was much more prominent in *D. mojavensis* than *D. arizonae*, which was evident throughout the time course of our analysis. While AS was detected for hundreds of genes in *D. mojavensis* at both timepoints, few genes in *D. arizonae* were alternatively spliced in response to mating, especially at 45 min. Altogether, distinct patterns of DE, AS, and functional enrichment between *D. mojavensis* and *D. arizonae* demonstrate considerable divergence in the female transcriptomic response to mating despite relatively recent common ancestry. These results are consistent with a prior study showing divergence in the proteomic response to mating in *D. simulans* and *D. mauritiana*^9^. Collectively, these studies suggest that the female postmating molecular response evolves rapidly, which is consistent with predictions of the coevolutionary divergence model.

The coevolutionary divergence model further predicts that divergence in the postmating response between species leads to incompatible interactions in heterospecific crosses. Moreover, given independence in coevolutionary trajectories during the process of divergence, the phenotypic outcomes and/or molecular mechanisms of incompatibilities are not necessarily expected to be the same in reciprocal crosses. We found disruptions in the normal female postmating transcriptional response of both species following copulation with heterospecific males, including disruptions in the normal (conspecific) patterns of DE and AS. Furthermore, several functional categories enriched in conspecific matings were not enriched in the heterospecific cross, indicating that transcriptional disruptions could have important functional consequences. While it is not clear whether these changes are causally linked to incompatibilities causing PMPZ isolation, the overall pattern is consistent with the coevolutionary divergence model.

Transcriptional disruption included misexpression/splicing of genes that normally respond to mating in conspecific crosses and genes that were not mating responsive in conspecific crosses but were differentially regulated following heterospecific copulation. This pattern was also observed in a previous transcriptional analysis of heterospecific mating in *D. mojavensis*^6^, and in a recent study of heterospecific matings between *D. novamexicana* and *D. americana*^7^. Misregulation of genes that normally respond to mating likely reflects disrupted interactions between male and female molecules in mismatched heterospecific crosses. While it makes sense that this transcriptional disruption could have adverse effects on reproductive outcomes, it is less clear to what extent the misregulation of non-mating responsive genes might also be involved in reproductive incompatibilities that give rise to PMPZ isolation. Interestingly, Ahmed-Braimah^7^ found a significant portion of such genes were linked to immunity, with heterospecific matings inducing a stronger immune response in females than conspecific matings. Similarly, we found enrichment of differentially regulated immune genes (humoral response) in heterospecifically mated *D. mojavensis* females. This cluster included four upregulated genes and two downregulated genes. Although enrichment analysis revealed overrepresentation of immune genes in *D. mojavensis* females only in the heterospecific cross, five of the six genes in the cluster were also differentially expressed in the same direction in the *D. mojavensis* conspecific cross and in both crosses involving *D. arizonae* females (though enriched only in the conspecific cross). Based on these data, we conclude that although mating induces differential regulation of several immune genes, there is not strong evidence that the immune response is heightened in females mated to heterospecific males relative to those mated to conspecifics. In general, differential regulation of immune related genes following mating is consistent with other studies in *Drosophila*, which have shown upregulation of immunity in response to both con- and heterospecific matings^7,40–45^. While we did find that most differentially expressed immune genes, including antimicrobial peptides (AMPs), were upregulated following mating, there were a few immune related genes that were downregulated. Most notably, *hemolectin* (*hml*), which is known to play a role in hemolymph clotting, was downregulated in all crosses. This is potentially relevant to the formation and/or degradation of the insemination reaction in the female reproductive tract of *D. mojavensis* and *D. arizonae* following mating. This mass at least superficially resembles a clot, and while it forms following both con- and heterospecific copulations, it is not efficiently degraded in the reproductive tracts of heterospecifically *D. mojavensis* females, contributing to PMPZ isolation^12^. The severity and time-course of the insemination reaction has not been examined in heterospecifically mated *D. arizonae* females.

Although gene expression and alternative splicing were disrupted in both heterospecific crosses, patterns of disruption were largely specific to each cross. For example, the proportion of genes with disrupted expression profiles was much lower for those that were mating-responsive in both conspecific crosses than for those that responded to mating in only one species (Fig 5). Moreover, the timing of and magnitude of transcriptional disruptions varied between the crosses, and the disrupted response in *D. arizonae* involved more genes that were only differentially regulated in the heterospecific cross compared to *D. mojavensis* (see Figs 2, 4 and 5). Together, these findings are consistent with the prediction that the underlying mechanisms of reproductive incompatibilities are likely to be different in reciprocal crosses as a result of independent coevolutionary trajectories.

The coevolutionary divergence model predicts that rapid evolution of traits involved in postcopulatory interactions leads to reproductive mismatches between males and females from different populations. In a transcriptomic context, this leads to the prediction that genes with more divergent transcriptional profiles in conspecific crosses should also display more disrupted profiles in heterospecific crosses. We found an overall positive correlation between conspecific transcriptional divergence and heterospecific disruption for both species across all timepoints. However, the magnitude and/or direction of the correlation differed for genes that were mating responsive in conspecific crosses versus those that were differentially regulated only in heterospecific crosses. Specifically, conspecific mating responsive genes showed a significantly positive correlation between divergence and disruption at all timepoints. In contrast, genes that were misregulated only in heterospecific crosses displayed either non-significant correlations or a weak negative correlation between divergence and disruption. Genes differentially expressed only as a result of heterospecific mating are by definition not a significant part of the normal mating (*i*.*e*. conspecific) response, and hence their evolutionary trajectories are not directly predicted by the coevolutionary divergence model. While we cannot directly assess whether transcriptional divergence in mating-responsive genes is driven by sexual selection and/or sexual conflict as predicted by the model, the difference in the relationship between transcriptional divergence and heterospecific disruption in mating vs. non-mating responsive genes suggests that different evolutionary forces may have acted on these sets of genes.

Although misregulated mating responsive genes tended to be more transcriptionally divergent in comparisons of conspecific crosses, we did not find evidence that misregulated genes, as a group, evolve rapidly at the protein coding sequence level as assessed by ω (Fig 8). In fact, all categories of misregulated genes evolved at similar rates to the genome or, in some cases, below the genome median. This aligns with the results of our earlier study, which also found that misregulated genes in heterospecific crosses between *D. mojavensis* females and *D. arizonae* males did not evolve more rapidly than other female reproductive genes^10^. Ahmed-Braimah^7^ also reported a similar result in crosses between *D. americana* and *D. novamexicana*, suggesting that this might be a general pattern in *Drosophila*. One explanation for this finding could be that rapid changes in protein structure are likely to be most consequential for male and female proteins that directly interact, which is consistent with previous studies showing rapid evolution of some female reproductive genes^45–50^. It is thus possible that many genes with disrupted expression profiles in heterospecific crosses are “downstream” of direct interactions between male and female molecules. As such, they may exhibit disrupted transcriptional profiles, while not evolving rapidly at the protein sequence level.

Overall, our data reveal that postmating transcriptional responses have diverged between *D. mojavensis* and *D. arizonae*, including striking changes in DE and AS that have potential functional implications. Moreover, we found that the normal transcriptomic response in each species was highly disrupted in heterospecifically mated females, with most mating responsive genes being misregulated in addition to genes that were only differentially regulated in heterospecific crosses. The patterns of disruption differed between the crosses, indicating potentially different mechanisms underlying reproductive incompatibilities. Importantly, mating responsive genes with more divergent transcriptional profiles in the two species also displayed more significant disruption in heterospecific crosses, as would be expected if transcriptional disruption reflects failed or suboptimal interactions between the female reproductive tract and components of heterospecific ejaculates. While these findings are consistent with predictions of the coevolutionary divergence model, we acknowledge that transcriptional disruption observed in heterospecific crosses may not be directly involved in incompatibilities that give rise to PMPZ isolation. Moreover, although transcriptional divergence between species and the level of transcriptional disruption in heterospecific crosses was positively correlated, the evolutionary forces that drove this divergence are not clear. Future research aimed at identifying specific male and female gene products that are directly involved in postmating incompatibilities is necessary to make causal links to PMPZ isolation and to test whether these genes have evolved under divergent coevolutionary trajectories of sexual selection and sexual conflict.

## Acknowledgements

We would like to thank undergraduate students Nathaniel Talamantes, Rorie Shae Robinson, Kamal Jitendra Patel, Moruj Athma and Graham Wegner from the University of Arizona for assistance in collecting the *Drosophila* samples. The research presented in this publication was supported by the Eunice Kennedy Shriver National Institute of Child Health and Human Development of the National Institutes of Health under award number R21HD097545 to LMM and JMB as well as by funds from the University of Arizona to LMM.

## Supplementary material

**Fig. S1.**
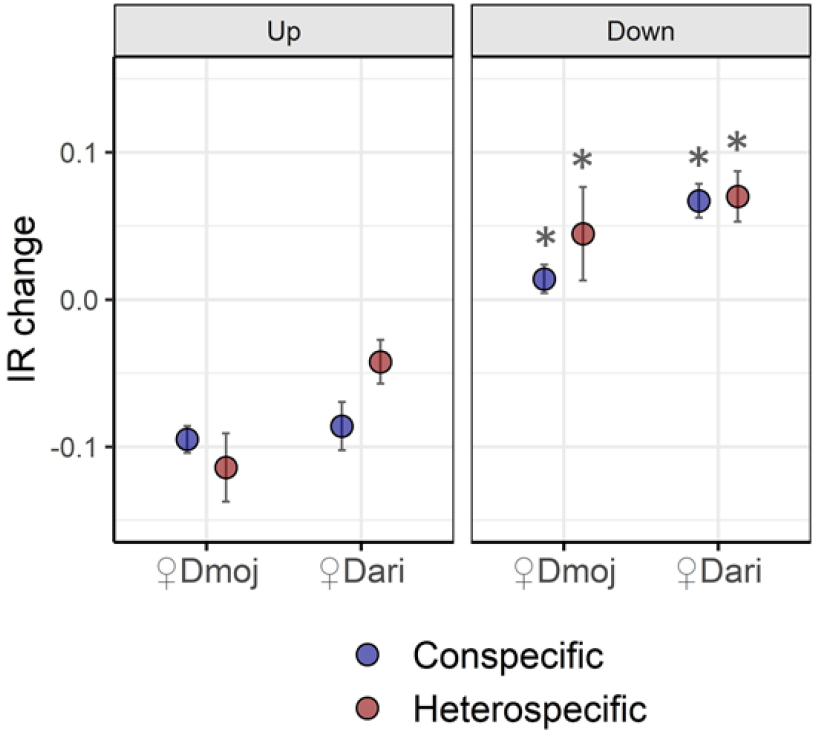
Changes in intron retention rates as estimated for up- vs down-regulated DE genes detected for con- and heterospecific matings between *D. mojavensis* and *D. arizonae*. *IR* change was estimated as *IR* mated – *IR* virgin samples. All mating experiments showed significant increase in of *IR* rates for down-regulated genes with respect to that of up-regulated ones. All significant comparisons with α = 0.05 following *GLM* analysis are indicated with * in the “down” plot. The *GLM* analysis was performed using categories of up and down regulation as independent variables and the level of IR change as the dependent variable for each mating experiment. *GLM* analysis was performed after square root transformation while accounting for normal distribution and homoscedasticity of the data.

**Fig. S2.**
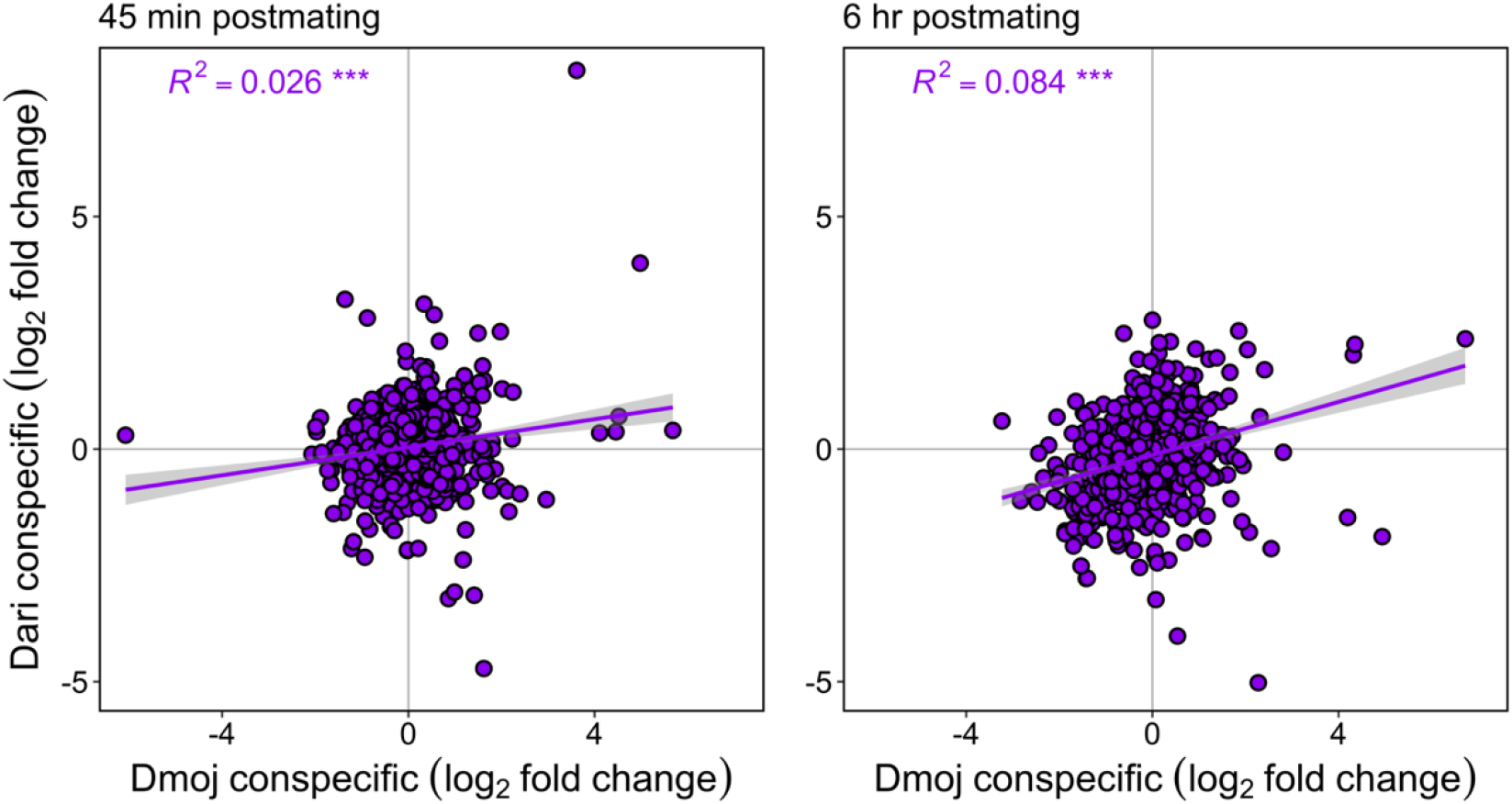
Patterns of transcriptional correlations between non-conspecific-responsive DE genes in *D. mojavensis* vs *D. arizonae*. All comparisons are performed against virgin females. Scatterplots indicate the expression change (log_2_) of non-significant DE genes (*FDR*_*α*_ > 0.05) for both species at 45 min postmating and 6 hrs postmating. Pearson’s *R*^*2*^ correlation coefficients and LM trend-lines (with 95% confidence intervals) between the species are indicated. P-values of correlations are noted: *** P < 0.001.

## Notes

### Competing Interest Statement

The authors have declared no competing interest.

